# Beta bursting in the retrosplenial cortex is a neurophysiological correlate of environmental novelty which is disrupted in a mouse model of Alzheimer’s disease

**DOI:** 10.1101/2021.04.26.441462

**Authors:** Callum Walsh, Thomas Ridler, Maria Garcia Garrido, Jonathan Witton, Andrew D. Randall, Jonathan T. Brown

## Abstract

The retrosplenial cortex (RSC) plays a significant role in spatial learning and memory, and is functionally disrupted in the early stages of Alzheimer’s disease. In order to investigate neurophysiological correlates of spatial learning and memory in this region we employed *in vivo* electrophysiology in awake, behaving mice, comparing neural activity between wild-type and J20 mice, a mouse model of Alzheimer’s disease-associated amyloidopathy. To determine the response of the RSC to environmental novelty local field potentials were recorded while mice explored novel and familiar recording arenas. In familiar environments we detected short, phasic bursts of beta (20-30 Hz) oscillations (beta bursts) which arose at a low but steady rate. Exposure to a novel environment rapidly initiated a dramatic increase in the rate, size and duration of beta bursts. Additionally, theta-beta cross-frequency coupling was significantly higher during novelty, and spiking of neurons in the RSC was significantly enhanced during beta bursts. Finally, aberrant beta bursting was seen in J20 mice, including increased beta bursting during novelty and familiarity, yet a loss of coupling between beta bursts and spiking activity. These findings, support the concept that beta bursting may be responsible for the activation and reactivation of neuronal ensembles underpinning the formation and maintenance of cortical representations, and that disruptions to this activity in J20 mice may underlie cognitive impairments seen in these animals.

## Introduction

The retrosplenial cortex (RSC) is considered to play a critical role in spatial learning and memory. Damage to this region results in severe impairments in navigation and landmark processing (see Mitchell *et al*., 2018 for review). There is a large body of experimental evidence suggesting the retrosplenial cortex is involved in the encoding, retrieval and consolidation of spatial and contextual memory (see Todd and Bucci, 2015 for review). Optogenetic stimulation of RSC neurons is sufficient to elicit retrieval and consolidation of contextual memories (Cowansage et al., 2014; De Sousa et al., 2019). RSC neurons encode a range of contextual information during navigation (Koike et al., 2017), and inactivation of the RSC during impairs performance in spatial memory and contextual fear memory tasks (Czajkowski et al., 2014; Kwapis et al., 2015), suggesting the RSC is involved in the storage of spatial information. Finally, Iaria *et al*., (2007) demonstrated that while hippocampal subregions are differentially involved in the encoding and retrieval of spatial information, the entire RSC is active during both processes. Spatial learning and memory impairments have been shown to be one of the earliest aspects of cognitive impairment in Alzheimer’s disease (AD). Patients exhibit disturbances in specific spatial memory processes associated with the RSC (Laczó et al., 2009; Vann et al., 2009; Morganti et al., 2013). During the early stages of AD, the retrosplenial gyrus has been shown to exhibit regional hypometabolism (as measured by FDG-PET), and considerable atrophy (Minoshima et al., 1997; Choo et al., 2010). As such, the RSC is a region of great interest in research into the brain’s function in health and AD.

Measurable correlates of brain function can have great value in fundamental neuroscience. They can aid the understanding of the complex ways in which the brain processes information and performs its many tasks, and also indicate how such functionality may be affected in disease. Similarly, these “functional biomarkers” can provide measurable benchmarks against which to test interventions which may affect or restore normal brain function (Walsh et al., 2017). Of the growing number of methodologies available for investigating brain function, *in vivo* electrophysiology remains a powerful tool with a superior temporal resolution to all others. The coordinated firing of large groups of neurons in the brain gives rise to waves of electrical activity, known as neural oscillations, which can be recorded as intracranial local field potentials (LFPs) or extracranial electroencephalograms (EEGs). It is thought that one of the roles of these oscillations in the brain is to coordinate the spiking activity of neurons, allowing computation and communication between potentially distant brain regions (Canolty et al., 2010). The temporal resolution of electrophysiology combined with the spatial specificity afforded by intracranial recordings make *in vivo* electrophysiology an invaluable tool for discovering functional correlates of brain function and disease-associated dysfunction.

In order to investigate the function of the RSC in spatial learning and memory, we recorded LFPs and multi-unit spiking activity from this region, while mice freely explored either a novel or familiar environment. To probe the effects of AD-associated amyloid pathology on RSC function we used J20 mice, a widely employed mouse model of amyloidopathy. In this manuscript, we describe short, phasic bursts of beta (20-30 Hz) oscillations, termed “beta bursts”, that occur within the RSC, while mice freely explore an environment. Beta bursting activity is significantly increased during exposure to a novel environment, and these bursts are correlated with increased neuronal spiking in the RSC. These data demonstrate that beta bursting in the RSC is a robust neurophysiological correlate of environmental novelty and may have a role in the storage and retrieval of cortical spatial representations. Finally, we observed aberrant beta bursting activity and an uncoupling of beta bursting from neuronal spiking in the RSC in J20 mice, which may disrupt its function, and underlie spatial learning and memory deficits seen in these mice (Cheng et al., 2007).

## Methods

### Ethics

All procedures were carried out in accordance with the UK Animal (Scientific Procedures) Act 1986 and were approved by the University of Exeter Animal Welfare and Ethical Review Body.

### Animals

8 male J20 mice and 6 wild-type littermates were bred at the University of Exeter and housed on a 12 hour light/dark cycle. Access to food and water was provided ad libitum. All mice underwent surgery at between 6-8 months of age. Mice were group housed prior to surgery, and single housed post-surgery, in order to prevent damage to the surgical implants.

### Surgery

Mice were unilaterally implanted with a 16 channel, single shank silicon probe (NeuroNexus Technologies, A1×16-5mm-100-177-CM16LP), in the right retrosplenial cortex (AP –2 mm, ML +0.5 mm, DV +1.75 mm, 0° Pitch). Mice were anaesthetised using isoflurane and fixed into a stereotaxic frame. A small craniotomy was drilled over the desired co-ordinate, and at least one hole was drilled in each of the major skull plates, in which miniature screws were placed to act as supports (Antrin Miniature Specialties). The probe was slowly lowered into the desired location, and fixed in place with dental cement (RelyX Unicem, 3M). The ground wire from the probe was connected to a silver wire, attached to a screw overlying the cerebellum. Throughout surgery, body temperature was monitored with a rectal probe and regulated by a feedback-controlled heat mat. Animals were kept hydrated by subcutaneous injections of Hartmann’s solution once per hour of surgery (0.01 ml/g body weight).

### Behaviour

After at least one week of post-surgical recovery, animals underwent a Novel/Familiar environment task, as shown in (Fig. 1). Individual mice were tethered to the recording apparatus, and placed in one of two high-sided recording arenas: one square, with black and white stripes, and one circular and lacking stripes. Both arenas each had two internal visual cues, placed on opposite sides. The animals were allowed to freely explore their environment for 15 minutes, after which, they were returned to their home cage. After 15 minutes in their home cage, the animal was returned to the same recording arena for another 15 minutes, and allowed to freely explore. Following this, the animal was returned to its home cage. This protocol was repeated at the same time of day, for 5 consecutive days, but on the fifth day, the animal was placed in the other, previously unseen arena. The order of exposure to these arenas was counterbalanced between animals. Each session can therefore be described by the experimental day, and the particular session within that day, with session A being the first, and session B being the second. Using this nomenclature, Sessions 1a and 5a were ‘novel’ sessions, while the remaining sessions were ‘familiar’ sessions. In order to reduce the stress associated with the recording process, animals were acclimatised to this process during a 10 minute test session 3 days prior to the start of the experiment, in which the animal was tethered and recorded from while in its home cage. An added benefit of this was to familiarize the animals with this experimental procedure, thus ensuring that perceived novelty during the first experimental session was limited to the environment, and not the experience of being tethered to the recording apparatus.

**Figure 1.**
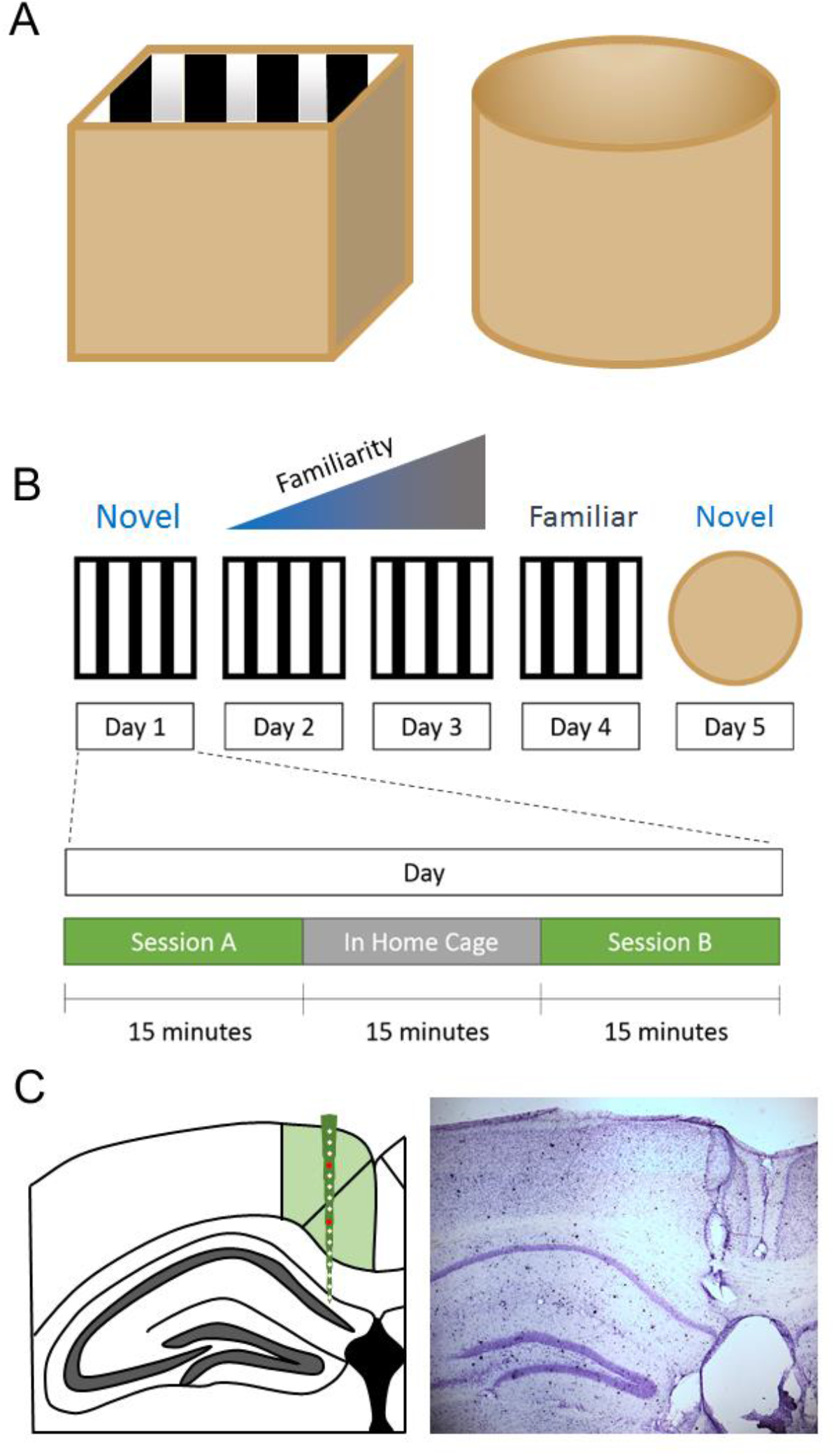
Experimental Design A. Diagrams of the recording arenas used for this study. Both are roughly equal sized, one is square, with black and white stripes along the walls and floor (left) and the other is cylindrical with plain brown floor and walls. B. Experimental procedure for the novel/familiar environment task. A mouse is placed in one of the recording arenas for two 15 minute sessions, referred to as sessions A and B, with a 15 minute break in their home cage between the two sessions. This is repeated in the same arena for 4 consecutive days, after which the arena is switched for the 5^th^ and final day. C. Single shank, 16 channel silicon probe electrodes were implanted in the retrosplenial cortex (green), so that they spanned the dysgranular (upper green section) and granular (lower green section) subregions. In order to verify the location of the electrodes, electrolytic lesions were made prior to perfusion, and slices were histologically prepared using Cresyl Violet stain. An example is shown (right).

### Data Collection

Throughout experimental sessions, Local Field Potentials (LFPs) were recorded using an Open Ephys Acquisition board (Open Ephys), which was tethered to the probe via a headstage (RHD 16-Channel Recording Headstage, Intan Technologies), and SPI cables (Intan Technologies). LFPs on each channel were sampled at 30 kHz, while the animal’s location was monitored using a pair of light-emitting diodes (LED) soldered to the headstage, and a video camera, placed directly above the arena. The location of these LEDs was tracked using Bonsai tracking software, so the location and running speed of the animal could be estimated offline.

### Data Analysis

LFPs were down-sampled (Spectral Analysis: 1 kHz, Burst Detection and Phase Amplitude Coupling: 3 kHz, Multi-Unit Activity: N/A) and de-trended, in order to remove any slow linear drift of the baseline that may occur across the session. The Chronux toolbox (Mitra and Bokil, 2008, http://chronux.org/) was used for the mtspecgramc function, as well as a number of built in MATLAB functions. All scripts used in this study were written in house, and are now publicly available (see Software Accessibility). All LFP analyses were performed for a single channel in the dysgranular and a separate single channel in granular RSC, except for multi-unit activity analysis, in which all channels in each region were used. The location of each channel was estimated from post-hoc histology.

### Power Spectra

Multi-taper spectral analysis was performed using the mtspecgramc function from the Chronux Toolbox, with a time-bandwidth product of 2 (1 second x 2 Hz), and 3 tapers, resulting in some smoothing of resulting spectra. The mtspecgramc function generates a power spectrogram by generating multiple power spectra for short segments of time series data, using a moving window; in our case with the window size of 1 s with no overlap. These spectrograms were then logged to the base 10, and multiplied by 10, in order to correct for the tendency of spectral power to decrease with a 1/f distribution. These individual spectra were averaged, resulting in a single mean power spectrum for the entire session, or for the first minute of each session, as specified in the results. Spectral data from 48 to 52 Hz, which incorporates line frequency noise (50 Hz), were removed, and linearly interpolated. The power of each frequency band was calculated as the mean power in each of the following frequency ranges: delta (1-5 Hz), theta (5-12 Hz), alpha (12-20 Hz), beta (20-30 Hz), low gamma (30-65 Hz), and high gamma (65-120 Hz).

### Beta Burst Detection

The data were band-pass filtered between 20-30 Hz, to isolate the beta frequency band. The amplitude and phase of this beta signal were calculated as the real and imaginary components of the Hilbert transform, respectively. The amplitude was z-scored, in order to give the instantaneous standard deviation of the beta signal amplitude from the mean. Epochs of the signal where this z-score exceeded 2 standard deviations from the mean amplitude were detected, as were the corresponding “edges” of these epochs, where the signal magnitude surpassed 1 standard deviation either side of the 2 standard deviation threshold. This was done in order to capture the time-course of these high beta amplitude epochs. Events that did not persist longer than a minimum duration of 150 ms (i.e. fewer than 3 oscillation cycles) were discarded. Furthermore, due to the sensitivity of this method to large, amplitude noise artefacts, any event whose peak amplitude exceeded three scaled median absolute deviations from the median of the events detected in that session were discarded as well. These remaining events were then considered beta-bursts. The duration and peak magnitude of each burst was calculated, as well as the distribution and total number of bursts in the session.

### Phase-Amplitude Coupling

To calculate phase-amplitude coupling, and create a comodulogram, modulation index was calculated individually for each pair of phase and amplitude frequencies. Modulation index was calculated as described by Canolty *et al*. (2006), with modification and vectorisation of some of the MATLAB code, for phase frequencies in bins of 0.25 Hz from 2 to 12 Hz, and for amplitude frequencies in bins of 2 Hz from 10 to 100 Hz. For each pair, local field potentials were filtered in the phase frequency band and the amplitude frequency band, after which the instantaneous phase and amplitude of each filtered signal was calculated, respectively, using the Hilbert transform. Subsequently, modulation index (MI) was calculated, but in order to attempt to reduce the possibility of spurious coupling, this was normalised through the use of 10 surrogates, created by time shifting the data by a random amount (between 1 and 59 seconds). In order to smooth the resulting comodulograms, the data matrix was linearly interpolated in both dimensions by a factor of 2.

### Multi-Unit Activity

Due to the distance between adjacent channels on the recording probe (100 µm) it is highly unlikely that activity of a single neuron would appear on multiple channels. Consequently, each channel was treated as an individual multi-unit. Raw local field potentials were first common average-referenced, using a mean of the signals from all other 15 channels, then filtered in the range of 500-14250 Hz, in order to isolate the spiking frequency band. Spikes were detected as peaks that crossed a threshold given by the median of the absolute voltage values of the signal, multiplied by 0.6745, as suggested by Quiroga, Nadasdy and Ben-Shaul (2004), and had a minimum separation of 0.5 ms. In order to investigate multi-unit activity during beta bursts, bursts were detected as previously mentioned, and bursts that occurred within a second of each other were discarded, to remove overlapping segments. A single peri-burst histogram was created for each channel by taking the total number of spikes in 20 ms time bins from 1 second before burst onset, to 1 second after, for all beta bursts. Each histogram was then normalised by dividing the count in each bin by the total number of spikes in all bins, averaged across all channels within the region, and then across all sessions, smoothed with a 100 ms moving mean filter, and z-scored with respect to the baseline epoch (1 second pre-burst).

### Software Accessibility

All code has been made publicly available at https://github.com/cfle/In-Vivo-Ephys-Code. This code is freely accessible for viewing, or use. If using any of this code in a paper, please, cite this paper as well as the GitHub repository (https://github.com/cfle/In-Vivo-Ephys-Code).

### Statistics

All statistical analysis was performed in MATLAB. Fourteen mice in total were used in this study, 6 wild-type and 8 J20, with each mouse undergoing a total of ten recording sessions (5 days, 2 sessions per day). Unfortunately, the local field potential data from Day 3 session 1 (i.e. session 3a) was corrupted for a single wild-type mouse, and therefore data for this mouse from this session was omitted from all figure making and statistics. Therefore the n numbers for all statistics are (wild-type: n = 6 (except from Day3a where n = 5), J20: n = 8). All statistics, unless stated otherwise, were performed using a two-way ANOVA, with genotype (wild-type/J20) and novelty (novel/familiar) as factors. It is important to note that due to the experimental design of our Novel/Familiar environment task, there were multiple novel and familiar sessions (2 novel, and 8 familiar). All sessions were either classified as novel or familiar and analysed accordingly. Following a significant main effect or interaction, Bonferroni-corrected multiple comparisons was performed, to investigate pairwise differences between different levels of either factor.

### Histology

Upon completion of the experiments, mice were killed using an overdose of sodium pentobarbital (Euthetal), and an isolated stimulator was used to produce electrolytic lesions at the recording sites. Mice were then transcardially perfused with 40% paraformaldehyde (PFA), and their brains were extracted and stored in PFA for 24 hours, after which they were transferred to phosphate-buffered saline (PBS) prior to sectioning. Brains were sliced into 100 µm sagittal sections using a vibratome (Leica), and stained with Cresyl Violet. Digital pictures were taken using QCapture Pro 7 software (Qimaging), and electrode sites were verified by comparing the lesion sites in these photographs to The Allen Mouse Brain Atlas (https://mouse.brain-map.org/static/atlas). Due to the high channel count of these probes, as well as their linear geometry, it was possible to account for small differences in the depth of each probe by selecting channels of similar depths across different probes. This resulted in reduced variability between animals in a range of neurophysiological measures.

## Results

To investigate neurophysiological correlates of spatial learning and memory in the retrosplenial cortex (RSC), local field potentials were recorded from across the entire dorsoventral axis of the RSC, while animals underwent a novel/familiar environment task. The RSC is made up of two distinct subdivisions: dysgranular (RSCdg), and granular (RSCg). While these regions are strongly interconnected, the neuroanatomical connectivity of these regions has been shown to differ (van Groen and Wyss, 1992; Van Groen and Wyss, 2003a, 2003b), therefore it is possible that the functional neurophysiology may vary as well, especially during a behavioural paradigm such as this, where spatial learning and memory processes may be stimulated. Due to the anatomical positioning of these subdivisions in rodents, it is possible to record from both RSCdg and RSCg at once, using a single, vertical silicon probe (Figure 1C). Therefore for this study, our analyses were performed for both subdivisions. We found very little difference between the electrophysiological activities seen in the two subregions. Furthermore, any changes seen in J20 mice were generally common to both subregions, with marginally greater effects in RSCg. For the sake of conciseness, we have decided to only show the data from RSCg in this paper.

### Spectral Analysis

Local field potentials from RSCg show a clear peak in theta frequency band (5-12 Hz) throughout recording sessions (Fig. 2a). In order to investigate any changes in oscillatory activity in RSCg during environmental novelty, power spectral analysis was performed on the entire 15 minutes of each session. These power spectra were averaged across novel and familiar sessions for wild-type and J20 mice. Beta and low gamma power were significantly higher overall during novel sessions (Main Effect Novelty - Beta: F(1,135) = 16.4,p = 8.8e-5; Low Gamma: F(1,135) = 10.8, p = 0.001, 2-way ANOVA). Furthermore, while alpha, beta, low gamma and high gamma power were significantly higher overall in J20 mice (Main Effect Genotype – Alpha: F(1,135) = 21.4, p = 8.46e-6; Beta: F(1,135) = 253, p = 1.01e- 32; Low Gamma: F(1,135) = 43.3, p = 9.56e-10; High Gamma: F(1,135) = 14.4, p = 2.3e-4, 2-way ANOVA), delta and theta power were significantly lower (Main Effect Genotype - Delta: F(1,135) = 9.23, p = 0.03; Theta: F(1,135) = 7.92, p = 0.006, 2-way ANOVA). Beta power was significantly higher during novel sessions in J20 (Nov: 17.7 ± 0.18; Fam: 16.9 ± 0.09; p = 4.7e-4) but not wild-type mice (Nov: 15.1 ± 0.21; Fam: 14.7 ± 0.1; p = 0.4). Upon closer inspection of power spectrograms (Fig. 2a), it was clear that spectral activity changed within novel sessions. Power in the alpha, beta and low gamma range appeared to be higher in the first minute of the session and diminish over time. As exemplified in (Fig. 2c), transient epochs of high power in the 12-40 Hz range are seen throughout the early stages of the session. By performing the same power spectral analysis as before on only the first minute of each session, clear differences appeared between novel and familiar sessions. Alpha, beta and low gamma power were significantly higher overall during novel sessions (Main Effect Novelty - Alpha: (F(1,135) = 5.73, p = 0.02; Beta: F(1,135) = 75.7, p = 1.01e-14; Low Gamma: F(1,135) = 35.6, p = 1.98e-8, 2-way ANOVA). Furthermore, alpha, beta, low gamma and high gamma power were significantly higher overall in J20 mice (Main Effect Genotype - Alpha: F(1,135) = 40.9, p = 2.47e-9; Beta: F(1,135) = 132, p = 1.1e-21; Low Gamma: F(1,135) = 14.1, p = 2.52e-4; High Gamma: F(1,135) = 12.9, p = 4.65e-4, 2-way ANOVA). Beta and low gamma power were significantly higher in wild type (Beta: Nov: 17 ± 0.28; Fam: 15 ± 0.14; p = 5.47e-8; Low Gamma: Nov: 14.6 ± 0.26; Fam: 13.2 ± 0.13; p = 3.62e-5) and J20 mice (Beta: Nov: 19.2 ± 0.25; Fam: 17.5 ± 0.12; p = 3.59e-8; Low Gamma: Nov: 15.1 ± 0.23; Fam: 14.2 ± 0.11; p = 0.002).

**Figure 2.**
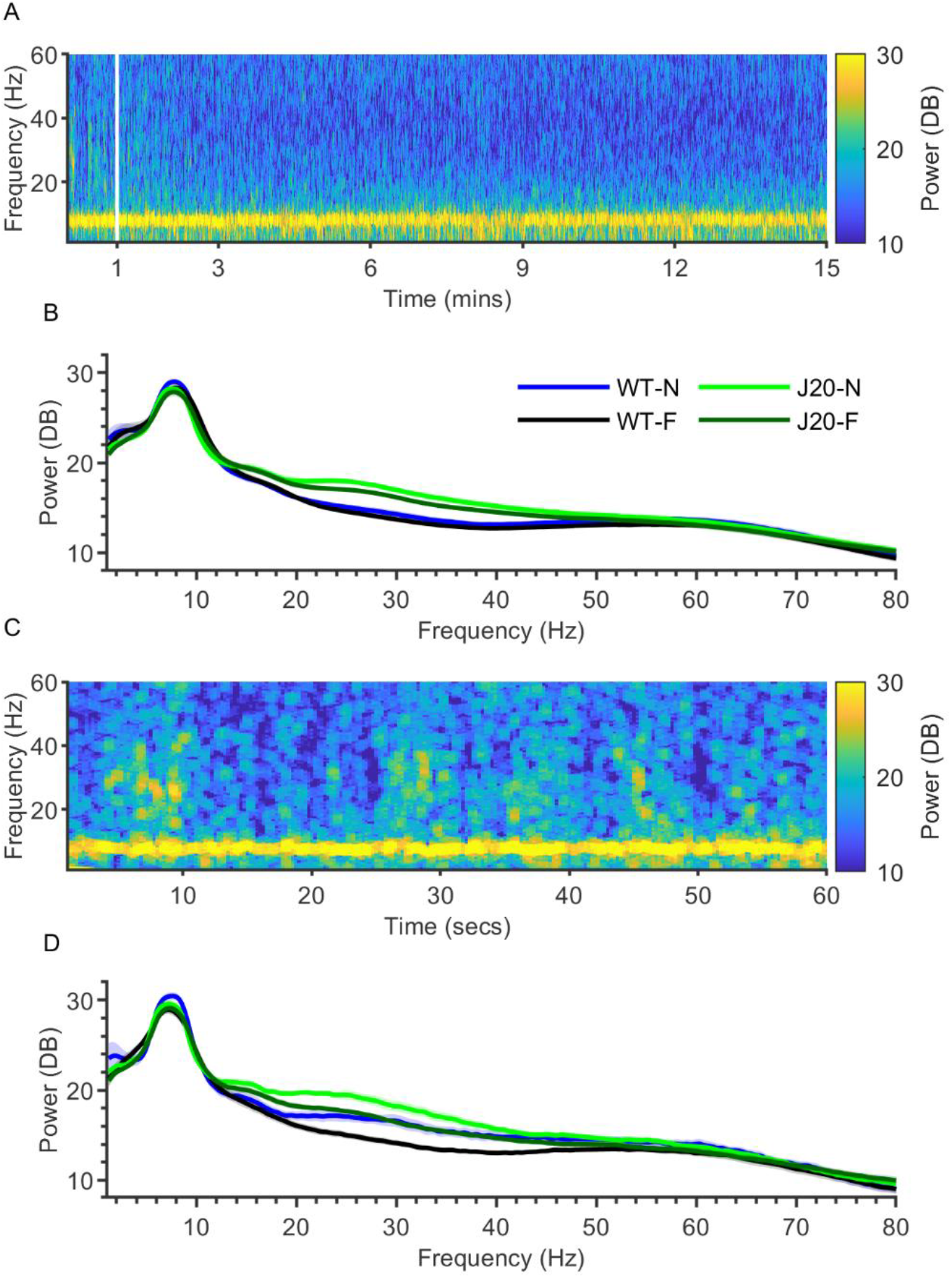
Beta (20-30 Hz) power is significantly higher during novelty in the granular retrosplenial cortex in wild-type and J20 mice. A. Example power spectrogram for an entire novel session in a wild-type mouse. B. Average power spectra for the entire 15 minutes of all novel and familiar sessions, for wild-type and J20 mice. Beta power was significantly higher during novelty in J20 (p = 4.7e-4) but not wild-type mice. When compared to WT power in the alpha, beta, low gamma and high gamma bands were significantly higher overall in J20 mice (p = 8.46e-6, p = 1.01e-32, p = 9.56e-10, 2.3e-4 respectively), whereas power in the delta and theta band were significantly lower (p = 0.03, p = 0.006 respectively). C. Example power spectrogram shown in A, expanded to show the first 60 seconds of the session. Short epochs of increased power in the 20-40 Hz range can be seen. D. Average power spectra for the first minute of all novel and familiar sessions, for wild-type and J20 mice. Beta and low-gamma power were significantly higher during novelty, for both wild-type (p = 5.47e-8, p = 3.62e-5 respectively) and J20 mice (p = 3.59e-8, p = 0.002 respectively). Alpha, beta, low gamma and high gamma power were significantly higher overall in J20 mice (p = 2.47e-9, p = 1.1e-21, p = 2.52e-4, p = 4.65e-4 respectively). (Data shown as mean ± SEM, WT: n = 6, J20: n = 8).

Across these time series, increased beta power occured in brief, discrete epochs, as shown in the expanded power spectrogram in (Fig. 3a). This can also be seen clearly in beta-filtered local field potentials, where these periods of high beta amplitude intersperse an otherwise very low amplitude oscillation. In order to understand the timescale and frequency domains of these events, wavelet analysis was used to investigate them further. As exemplified in (Fig. 3C), these individual events were short in duration, and peaked in the 20-30 Hz, beta band.

**Figure 3.**
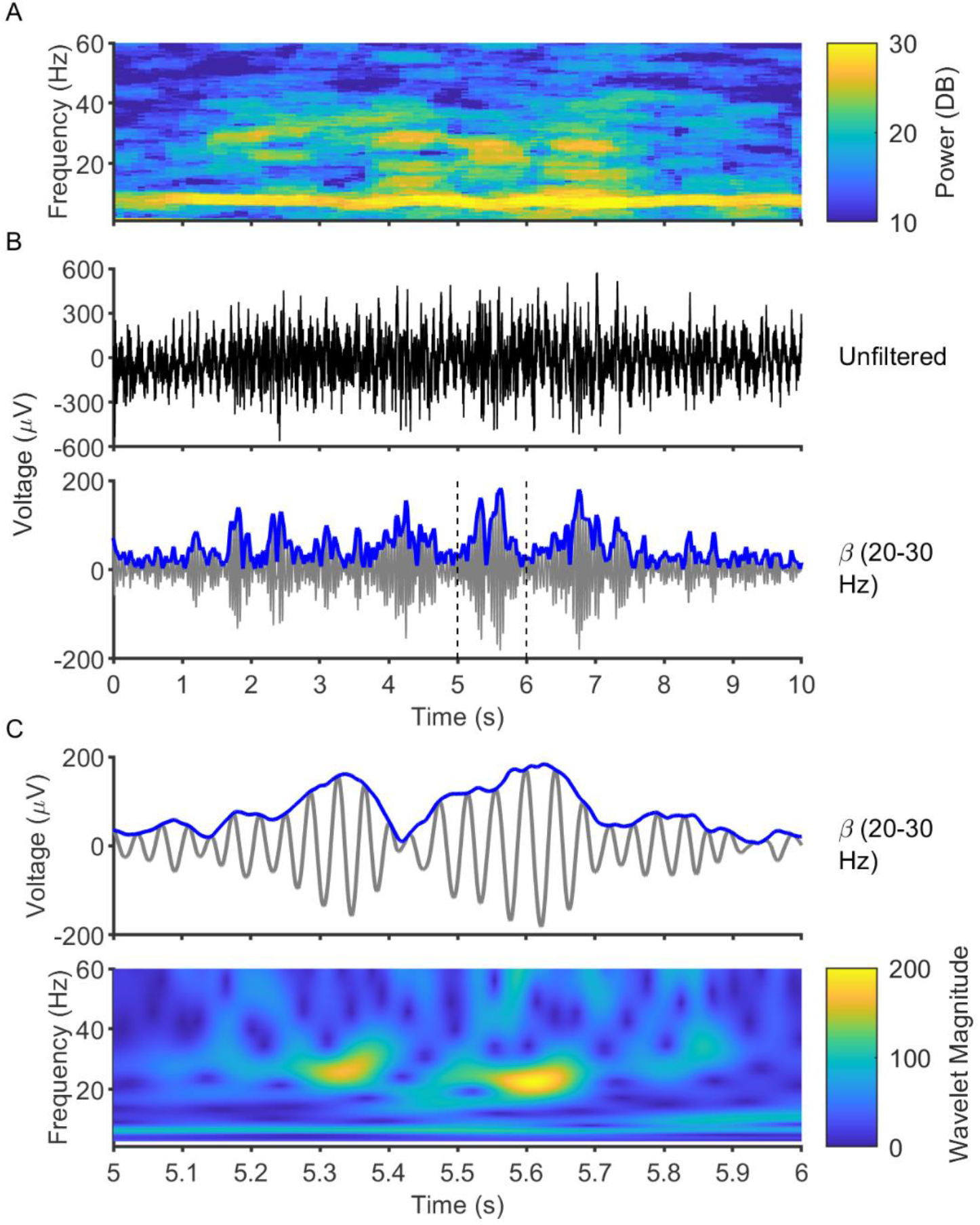
Retrosplenial local field potentials are marked by short, phasic increases in beta power, referred to as beta bursts. A. Example power spectrogram showing transient increases in beta power. B. Local field potentials of data shown in A, both unfiltered (top), and filtered in the beta band (bottom), with the envelope amplitude in blue for clarity. The beta-filtered local field potential shows clear epochs of high beta amplitude, which intersperse a low amplitude continuous beta oscillation. C. Expanded trace of the dashed area in shown in B (top), and a continuous wavelet spectrogram of this time series (bottom). Due to the high temporal resolution of wavelet-based methods, these periods of high beta amplitude can be seen to be brief in duration, only lasting around 100-200ms.

### Beta Bursting Activity

In order to investigate this phasic beta activity in more depth, an algorithm was written to detect and analyse these “beta bursts”; the basis of this algorithm is illustrated in (Fig. 4a). Once all putative bursts have been detected, the duration and magnitude of these beta bursts was calculated (Fig. 4a). With these transient epochs of high beta power now classified as discrete beta bursts, it is possible to compare this beta activity between sessions. Overall, there were significantly more beta bursts detected during novel sessions compared to familiar sessions (Main Effect Novelty - F(1,135) = 74, p = 1.73e-14, 2-way ANOVA). As shown in (Fig. 4b), there were significantly more beta bursts detected during novelty, for wild-type (Nov: 33.7 ± 2.42; Fam: 21.4 ± 1.22; p = 7.59e-5) and J20 mice (Nov: 56.3 ± 2.1; Fam: 37.8 ± 1.05; p = 4.83e-12). Furthermore, on average the number of beta bursts detected was significantly higher in J20 mice (Main Effect Genotype – F(1,135) = 118, p = 3.45e-20, 2-way ANOVA). Furthermore, it is possible to investigate the distribution of beta bursts within sessions. As shown in (Fig. 4c), during familiar sessions the rate of beta busting was reasonably steady, as indicated by the linear relationship between time and burst number shown in the cumulative frequency plot, for both wild-type and J20 mice. During novel sessions, however, there was a high rate of beta bursting during the first 1-3 minutes of the session, which gradually decreased over time to a steady rate. The rate of beta bursting was significantly higher in J20 mice during familiar sessions, and during the initial and final part of novel sessions.

**Figure 4.**
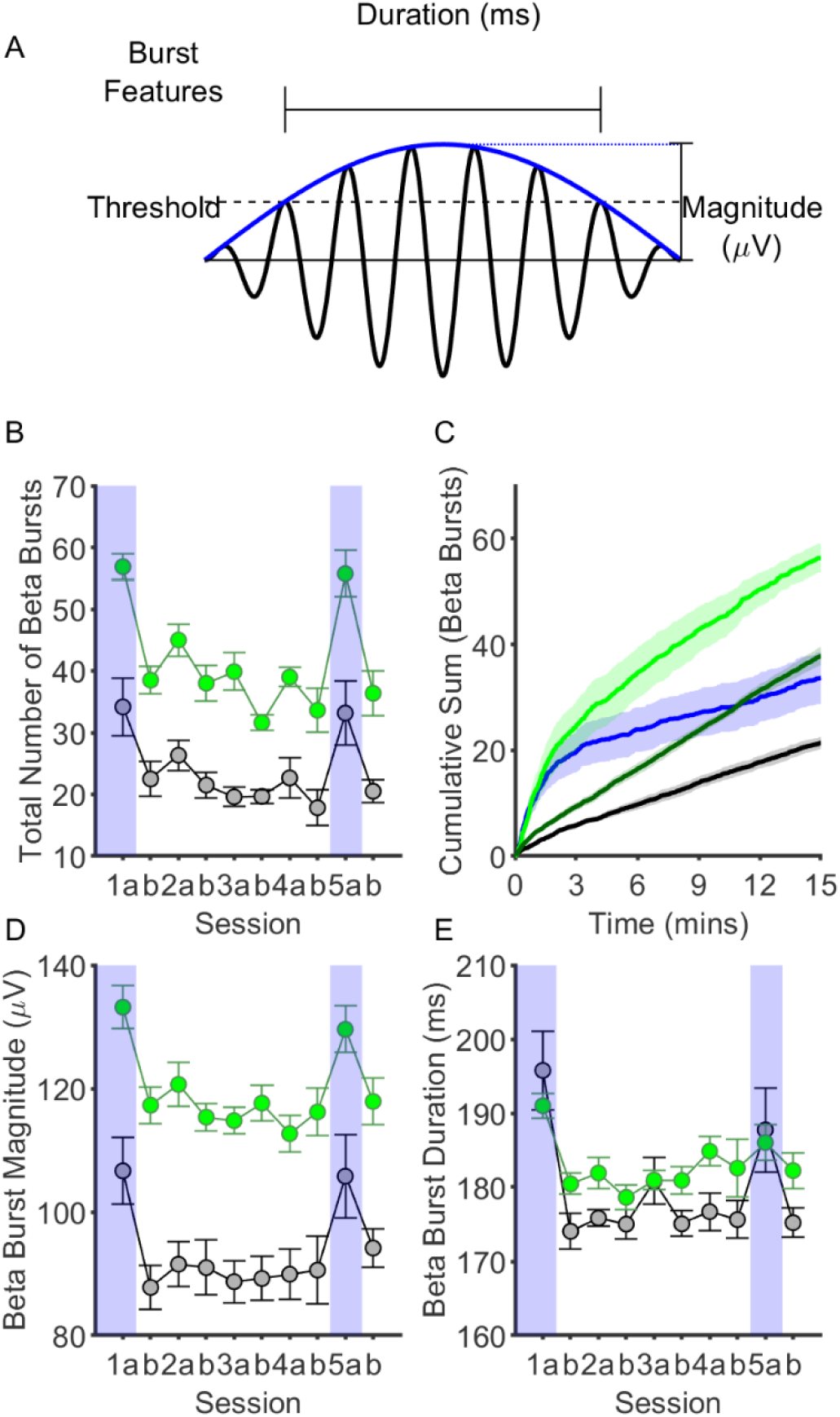
Beta bursting activity in the granular retrosplenial cortex (RSCg) is highly associated with novelty, and dysregulated in J20 mice. A. Diagram illustrating how beta bursts were detected, as well as how the magnitude and duration of these events were calculated. B. Graph showing the average number of beta bursts detected in RSCg in each session, for wild-type (black) and J20 mice (green). Novel sessions Day1a and Day5a are highlighted in blue for clarity. There were significantly more beta bursts in novel sessions as compared to familiar sessions, for both wild-type (p = 7.59e- 5) and J20 mice (p = 4.83e-12). C. Cumulative frequency graphs of number of bursts detected in novel and familiar sessions, for wild-type and J20 mice, showing the time course of bursting activity within sessions. While beta bursting occurred monotonically during familiar sessions, during the first 2-3 minutes of a novel session, beta bursting was substantially increased. D. Graph showing the average magnitude of beta bursts in RSCg in each session, for wild-type and J20 mice. Beta bursts were significantly larger in magnitude in novel sessions, for wild-type (p = 2.88e-5) and J20 mice (p = 5.16e-6). Beta bursts were also, on average, significantly larger in magnitude in J20 mice (p = 2.97e-22). E. Graph showing the average duration of beta bursts in RSCg in each session, for wild-type and J20 mice. Beta bursts were significantly longer in duration in novel sessions, for wild-type (p = 3.32e-9) and J20 mice (p = 0.005). (Data shown as mean ± SEM, WT: n = 6, J20: n = 8).

The features of these beta bursts may also vary depending on novelty and genotype. Burst magnitude was significantly higher overall during novel sessions (Main Effect Novelty - F(1,135) =48.7, p = 1.21e- 10, 2-way ANOVA). Furthermore, burst magnitude was significantly higher overall in J20 mice (Main Effect Genotype - F(1,135) = 137, p = 2.97e-22, 2-way ANOVA). As shown in (Fig, 4d), beta bursts were significantly larger in magnitude during novelty, for both wild-type (Nov: 106 ± 2.96; Fam: 90.4 ± 1.5; p = 2.88e-5) and J20 mice (Nov: 131 ± 2.57; Fam: 117 ± 1.28; p = 5.16e-6). There was also a significant interaction between the effects of genotype and novelty on beta burst duration (F(1,135) = 8.04, p = 0.005, 2-way ANOVA). As shown in (Fig.4e), beta bursts were significantly longer in duration during novel sessions for both wild-type (Nov: 192 ± 2.1; Fam: 176 ± 1.1; p =3.32e-9) and J20 mice (Nov: 189 ± 1.8; Fam: 182 ± 0.9; p = 0.005).

### Phase-amplitude Coupling

As elegantly shown by van Ede *et al*. (2018), continuous oscillations may appear as phasic burst events if their amplitude varies greatly over time. The amplitude of high frequency oscillations such as gamma may be modulated by the phase of low frequency oscillations such as theta (Canolty et al., 2006). This interaction is generally thought to allow slow, large amplitude oscillations to coordinate faster, small amplitude local oscillations. For this reason, it was of interest for us to investigate whether the amplitude of beta oscillations was coupled to the phase of theta oscillations, an increase in which may underlie the increased beta bursting activity seen during novelty. As shown in (Fig. 5a), phase-amplitude coupling efficacy was calculated for a range of phase and amplitude frequencies, and the effect of novelty and genotype determined. The strength of phase-amplitude coupling was quantified for theta-alpha, theta-beta and theta-gamma coupling for each session (Fig. 5b). There were significant interactions between the effects of genotype and novelty for theta-alpha coupling (F(1,135) = 12.8, p = 4.72e-4) and theta-beta coupling (F(1,135) = 17.7, p = 4.73e-5, 2-way ANOVA). Theta-alpha coupling was significantly higher in novel sessions for wild-type (Nov: 2.59 ± 0.15; Fam: 1.6 ± 0.07; p = 2.4e-7) but not J20 mice (Nov: 2.2 ± 0.13; Fam: 1.98 ± 0.06; p = 1). Theta-beta coupling was also significantly higher in novel sessions for wild-type (Nov: 1.65 ± 0.08; Fam: 1.16 ± 0.04; p = 1.04e-6) but not J20 mice (Nov: 1.23 ± 0.07; Fam: 1.23 ± 0.03; p = 1). There were no significant effects of novelty on theta-gamma coupling, but theta-gamma coupling was lower on average, in J20 mice (Main Effect Genotype – F(1,135) = 19.7, p = 1.87e-5). It is important to note that in order to focus on the most physiologically and behaviourally relevant part of the session, this analysis was performed for the first minute of each session. When the same analysis was performed on the last minute of each session, there was no effect of novelty on coupling in any band for either genotype (data not shown).

**Figure 5.**
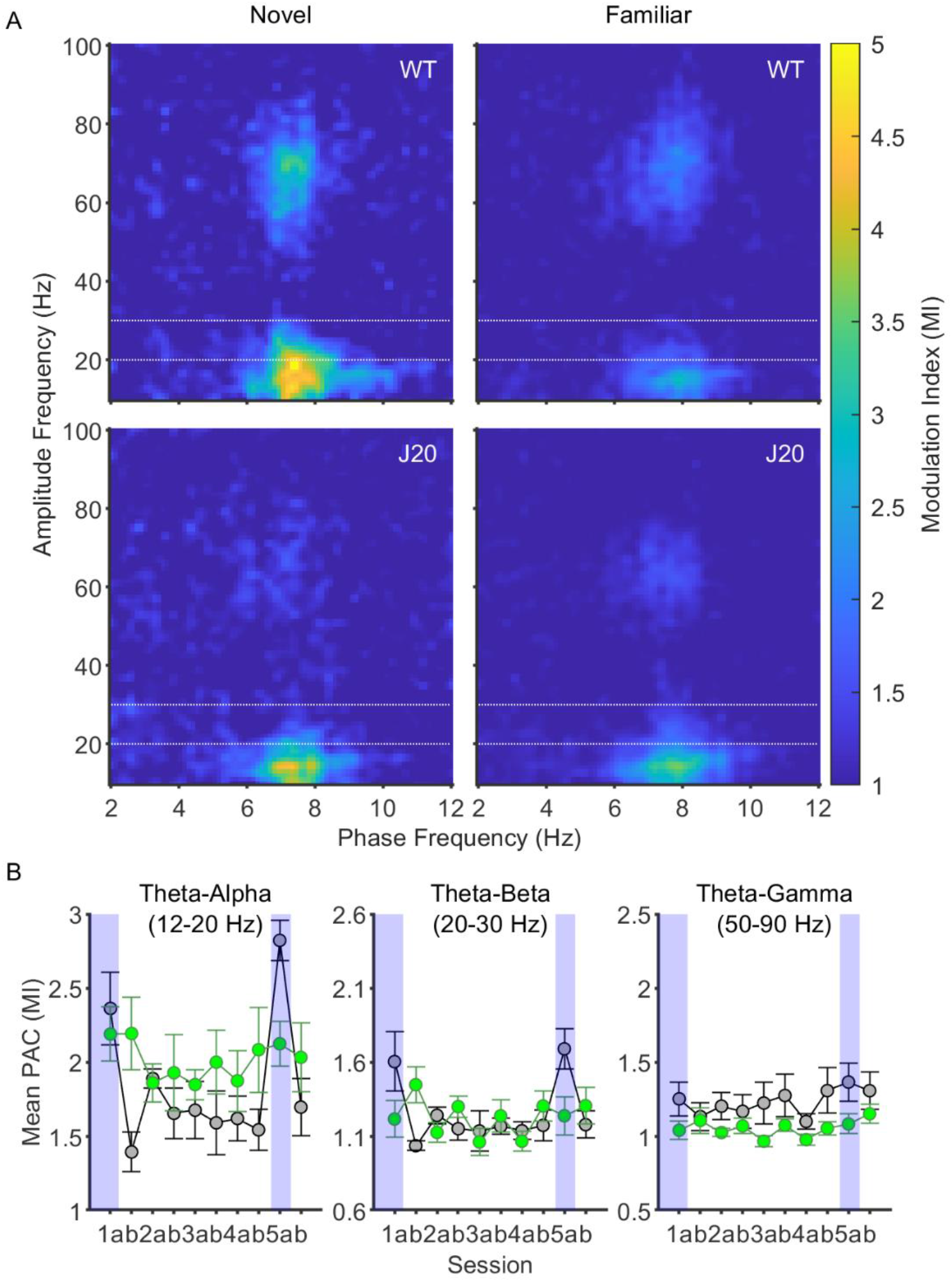
Theta-alpha and theta-beta phase-amplitude coupling are increased during novelty in the granular retrosplenial cortex (RSCg). A. Average comodulograms showing the strength of cross-frequency phase-amplitude coupling in RSCg during the first minute of novel and familiar sessions, for wild-type and J20 mice. Note the presence of three peaks in the first comodulogram, in the theta-alpha, theta-beta and theta-gamma ranges (the boundaries of which are denoted by the dotted lines). B. Average MI in the theta-alpha (left), theta-beta (center) and theta-gamma ranges (right), for each session, for wild-type (black) and J20 mice (green). Novel sessions Day1a and Day5a are highlighted in blue for clarity. Theta-alpha and theta-beta coupling were significantly higher in novel sessions for wild-type mice (p = 2.4e-7, p = 1.04e-6 respectively), but not J20 mice. (Data shown as mean ± SEM, WT: n = 6, J20: n = 8).

### Spiking Activity

In order to determine whether beta bursting was associated with a change in neuronal firing, multi-unit activity was investigated. Due to the linear geometry of the silicon probes, and the 100 µm distance between channels, it was not possible to reliably identify single unit activity, as activity from a single neuron was unlikely to appear on multiple channels, limiting spatiotemporal clustering methods such as those enabled by tetrodes or higher density silicon probes. Therefore, spikes appearing on a single channel could be from one or more nearby neurons. This, however, does mean that it is possible to treat each individual probe channel as a single multi-unit, to facilitate investigation of the relationship between neuronal spiking activity and beta bursting. As shown in the left panel of (Fig. 6a), individual spike waveforms can be readily discerned, and these spike waveforms are similar in wild-type (black) and J20 (green) mice. Furthermore, there was a trend towards higher multi-unit firing rate in J20 mice compared to wild-type mice (WT: 12.9 Hz ± 4.9; J20: 33.5 Hz ± 7.3; t(12) = -2.18, p = 0.05; unpaired t-test, Fig. 6a, right). The average beta amplitude during beta bursts is shown in (Fig. 6b), averaged across all bursts with non-overlapping time segments. Beta bursts in both genotypes are associated with a brief, monophasic increase in beta amplitude that lasts no more than 200 ms on average. Finally, (Fig. 6c) shows peri-event time histograms for spike rate during beta bursts, as a Z score from the pre-burst baseline (left of the dotted line). In order to investigate statistically significant changes in spike rate during bursts, the maximum z scored spike rate was determined at the peak of beta amplitude (approximately 100 ms after burst onset), for each animal, and compared to the mean pre-burst spike rate (0 due to z scoring of spike rate to baseline) using a one-sample t-test. Beta bursting in the RSCg of wild-type mice was associated with a significant increase in spike rate during beta bursts (Z-scored spike rate from baseline: 2.24 ± 0.46, t(5) = 4.86, p = 0.005, one-sample t-test; Figure 6c, left). Conversely there was no significant increase in spike rate during beta bursts in J20 mice (Z-scored spike rate from baseline: 0.78 ± 0.39, t(7) = 1.98, p = 0.09, one-sample t-test; Figure 6c, right). The difference between spike rate during beta bursts in wild-type and J20 mice, as determined by a two-sample t-test, was significant (t(12) = 2.4, p = 0.03, two-sample t-test). These data indicate that beta bursts are closely coupled to neuronal spiking in RSCg in wild-type mice, and that this relationship is effectively uncoupled in J20 mice.

**Figure 6.**
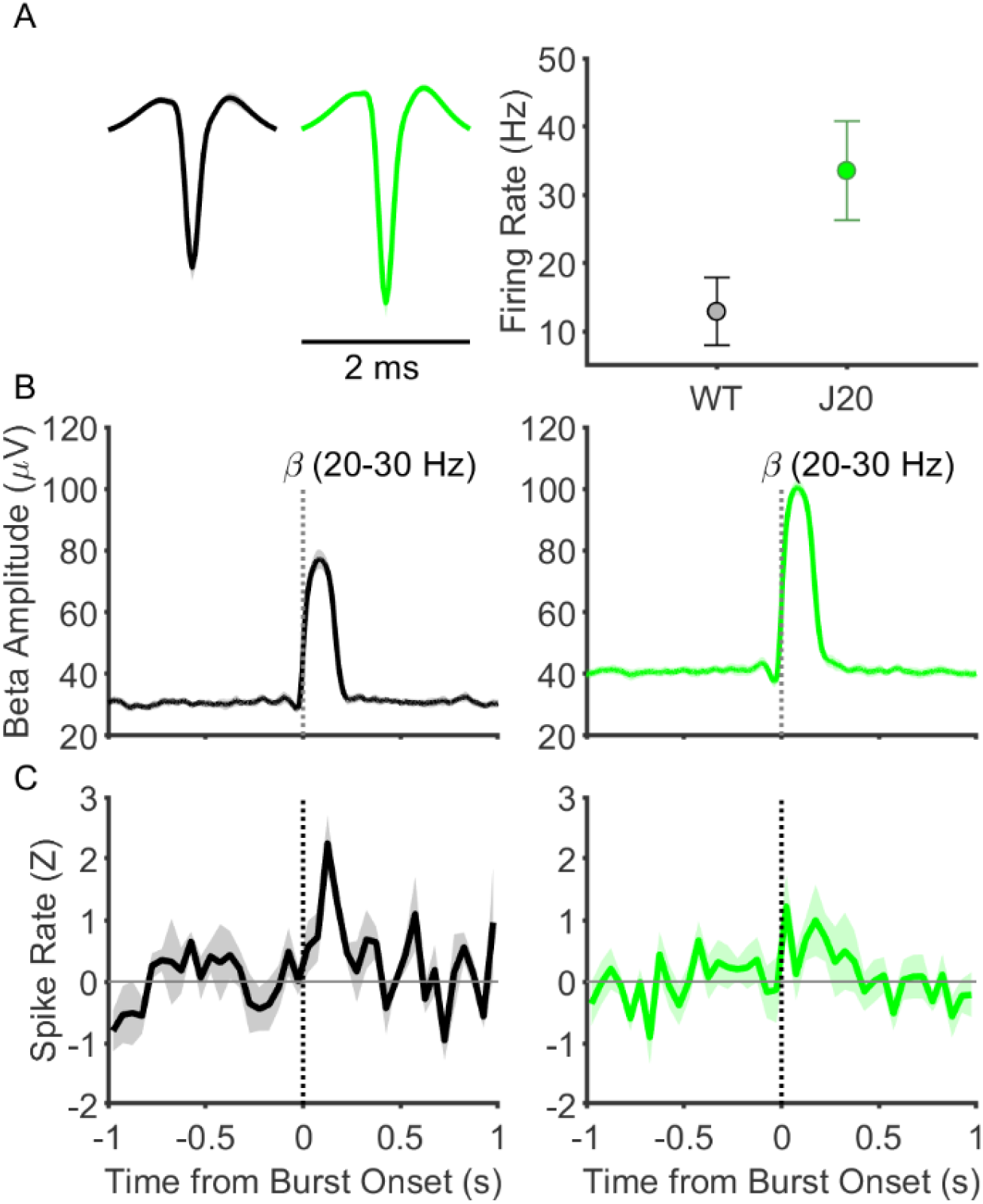
Spiking activity in RSCg is coupled to beta bursting in wild-type mice, but disrupted in J20 mice. A. Average spike waveforms for multi-unit activity in wild-type (black) and J20 (green) mice (left) and graph of average firing rate for detected multi-units across all sessions (right). There was a trend towards increased multi-unit firing rate in J20 mice compared to wild-type mice (p = 0.052, unpaired t-test). B. Graphs showing beta amplitude over time for beta bursts, time locked to the onset of the burst (dotted line), and averaged across all detected bursts, for wild-type mice (left) and J20 mice (right). Beta bursting was associated with a monophasic increase in beta amplitude that returns to baseline after around 250 ms. C. Peri-event histograms showing multi-unit activity spike rate during beta bursts, for wild-type (left) and J20 mice (right). Data is shown as Z score from baseline (pre-burst epoch), and averaged across all beta bursts with non-overlapping time segments. Dotted vertical line denotes the burst onset, while the solid horizontal line is shown to indicate the baseline of zero. Spike rate significantly increased during bursts in wild-type mice (p = 0.005), but not in J20 mice (p = 0.09).

## Discussion

In this study we attempted to identify neurophysiological correlates of environmental novelty in the mouse retrosplenial cortex (RSC), and investigate how these may be affected by amyloid pathology. We observed phasic increases in the amplitude of beta frequency neuronal oscillations, termed beta bursts, which occurred more frequently and with larger amplitude during novelty, and were positively correlated with neuronal spiking. A number of aberrant neurophysiological changes were seen in the RSC in J20 mice. Alpha, beta and low gamma power were significantly increased, and increases in beta bursting activity were seen during both novelty and familiarity. Beta bursts were more frequent, and larger in magnitude, yet the coupling of beta bursts to spiking activity was lost, suggesting a functional uncoupling of beta bursting with local neuronal activity. Finally, theta-beta phase-amplitude coupling was also disrupted, resulting in a loss of an effect of novelty on this activity. These results together indicate that beta bursting activity is a neurophysiological correlate of environmental novelty in the RSC, which is disrupted in J20 mice.

Numerous studies have noted changes in beta activity in a range of brain regions, during a variety of behaviours (see Spitzer and Haegens, 2017 for review). It is important to note that due to variability between groups in the naming and frequency ranges of neural oscillation frequency bands, cross-study comparison is often complicated. What we have referred to as beta, has previously been called upper beta (Spitzer and Haegens, 2017), beta2 (França et al., 2014), or slow gamma (Carr et al., 2012; Remondes and Wilson, 2015). For the sake of clarity, references to beta oscillations in this paper refer to the 20-30 Hz frequency range. Others have noticed similar novelty-induced beta oscillations in the hippocampus: Berke *et al*. (2008) reported a large increase in beta power that appeared when mice explored a novel environment, which persisted for around a minute, before returning to a lower level. The authors concluded that these oscillations may be a “dynamic state that facilitates the formation of unique contextual representations.” As shown in Igarashi *et al*. (2014), coherent 20-40 Hz oscillatory activity increased between the hippocampus and lateral entorhinal cortex during odour discrimination, and coincided with the development of odour-specific neural representations in these regions. Work by França *et al*. (2014) demonstrated that beta power was also transiently enhanced in the hippocampus during exploration of novel objects, but not previously experienced familiar items. Furthermore, they found that administration of an amnestic agent, namely haloperidol, resulted in a similar increased beta activity upon re-exposure to previously encountered objects, suggesting they had been “forgotten” and were therefore novel again. This further reinforces the idea that hippocampal beta activity is related to novelty, and extends the previous work by demonstrating that hippocampal-dependent novel object recognition can also elicit beta oscillations. Subsequently, França, Borgegius and Cohen (2020) investigated novelty-associated beta bursting in a larger hippocampal novelty circuit, by simultaneously recording from hippocampus, prefrontal cortex and parietal cortex during environmental and object novelty. Novelty-associated increases in beta power were seen in the prefrontal cortex during environmental novelty, and authors demonstrated significant phase-amplitude coupling of delta and theta to beta oscillations, which were increased in novelty. Similarly, in the RSC we see strong coupling between theta phase and beta amplitude, which is significantly higher during novelty, but only in wild-type mice. Others have noted theta-beta PAC in humans as well, both in the hippocampus during a working memory task (Axmacher et al., 2010), and in the inferior temporal cortex during object novelty (Daume et al., 2017). Interestingly, the studies mentioned above tend to view beta activity as continuous oscillations, rather than discrete events. This is despite Berke *et al*. (2008) noting that beta appears as pulses, and a brief mention of burst detection and characterisation by França *et al*. (2014). As demonstrated in this study, novelty-associated beta oscillations in the RSC conform well to a model of discrete bursts, where their rate, magnitude and duration can vary depending on environmental novelty. Due to the use of averaging across trials or analysis spanning long temporal segments, the phasic nature of transient oscillatory events can be easily lost. Furthermore, in the somatosensory cortex, beta synchronicity appears in short events in both mice and humans; the features of which, such as duration and frequency range, are highly conserved across tasks and species (Shin et al., 2017).

Beta oscillations have long been associated with motor activity and sensory processing, and a large body of work has also noted changes in beta activity in a range of brain regions during other cognitive tasks (see Engel and Fries, 2010 for review). This gave rise to the hypothesis that the unifying function of beta oscillatory activity in these different regions was the maintenance of the “status-quo”, be it the current motor state, sensory stimulus or cognitive set (Engel and Fries, 2010). This theory would suggest that, beta activity would be decreased during novelty, and increased during familiarity. As we have shown, this is not the case. While steady and persistent beta bursting during familiarity may support the maintenance of the contextual “status-quo”, in this case the environment, this theory does not reconcile the significant increases in beta activity that occur during novelty.

Many groups have previously shown that information may be rapidly represented and stored in the RSC (Cowansage et al., 2014; Czajkowski et al., 2014; Koike et al., 2017; Vedder et al., 2017). Beta oscillations have also been shown to carry a variety of different forms of contextual information in a range of brain regions, and phasic increases in beta power during working memory maintenance may represent reactivation of encoded information (Spitzer and Haegens, 2017). Supporting this is a study in which the authors employed transcranial magnetic stimulation to activate a currently unattended memory, as shown by an increase in content-specific beta activity (Rose et al., 2016). The theory put forth by Spitzer and Haegens (2017), is that beta oscillations can activate and reactivate neuronal ensembles to create and recall cortical representations. This theory is consistent with the data shown in this study: high beta bursting activity during perceived novelty activates neurons in the RSC, which may encode content about the novel environment, and subsequent beta bursting may continuously reactivate these ensembles, further consolidating or altering this representation. Recent breakthroughs in real-time burst detection and neurofeedback have made it possible to artificially induce beta bursts in awake behaving animals, creating the possibility of testing this hypothesis directly (Karvat et al., 2020).

A number of neurophysiological changes were seen in the RSC in J20 mice. Increases in alpha, beta and gamma power are indicative of a hyperexcitability phenotype, which has been previously noted in this strain (Palop et al., 2007; Palop and Mucke, 2009). Increases in beta bursting rate and burst magnitude were also notable. Finally, and most importantly, beta bursting activity was effectively uncoupled from neuronal spiking in J20 mice, potentially impairing the ability to form neuronal ensembles that encode and store information in the RSC. At the age point used, amyloid pathology in J20 mice is thought to be predominantly located in the hippocampus in this model, although, amyloid pathology seems to develop in the RSC to a much greater extent than other cortical regions, especially in RSCg (Whitesell et al., 2019). Hyperexcitability of cortical neurons in a mouse model of amyloid pathology was more prevalent in neurones proximal to amyloid plaques (Busche et al., 2008), and inhibitory interneuron dysfunction in J20 mice has been shown to lead to cortical network hypersynchrony and spontaneous epileptiform discharges (Verret et al., 2012). The hippocampus projects directly to RSCg, and indirectly, via the subiculum, to RSCdg (van Groen and Wyss, 1992; Van Groen and Wyss, 2003a, 2003b), so network dysfunction in RSC may be explained by its high levels of amyloid pathology or its anatomical connectivity with an increasingly dysfunctional hippocampus (Palop et al., 2007).

These findings demonstrate a novel form of Alzheimer’s disease (AD) related cortical dysfunction, which may underlie or exacerbate cognitive dysfunction seen in these mice, and in people with AD. Erroneous attribution of novelty to familiar environments, could cause memory impairments, and result in wandering and confusion. Interestingly, aberrant beta bursting has long been associated with another progressive neurodegenerative disease, Parkinson’s disease. Increased beta oscillatory activity in the basal ganglia and cortex are associated with motor impairments in Parkinson’s disease (for review see Brittain, Sharott and Brown, 2014), and administration of levodopa has been shown to improve motor function and reduce beta oscillations (Brown et al., 2001). The loss of coupling between beta bursting and neuronal spiking seen in J20 mice suggest that attenuating bursting without restoring this coupling may be ineffective in AD. Furthermore, the dysfunction in novelty-associated beta bursting identified in this study may be a useful functional biomarker of AD-related amyloidopathy, which could be used to measure the neurophysiological effectiveness of possible disease modifying therapeutics.

In conclusion, phasic bursts of beta oscillations may be a functional means of activating neural ensembles to form, and subsequently reactivate cortical representations. Network dysfunction in J20 mice results in aberrant beta bursting and an uncoupling of beta bursting from spiking, which may underlie cognitive impairments in these mice.

## Acknowledgements

CFW was funded by a University of Exeter and Janssen Pharmaceutica studentship. TR was supported by an ARUK Major Project grant (ARUK-PG2017B-7) awarded to JTB and AR.

## Notes

### Competing Interest Statement

The authors have declared no competing interest.

